# Induction of intercrypt goblet cells upon bacterial infection

**DOI:** 10.1101/2024.07.03.601821

**Authors:** Renaud Léonard, Ewa Pasquereau-Kotula, Edwige Madec, Benjamin Marsac, Adriana Mihalache, Laurence du Merle, Jordan Denis, Corentin Spriet, Philippe Sansonetti, Shaynoor Dramsi, Catherine Robbe Masselot

## Abstract

Intestinal mucins play a crucial role in the mucosal barrier, serving as the body’s initial defense against microorganisms. However, how the host regulates the secretion and glycosylation of these mucins in response to bacterial invasion remains unclear. Our study demonstrates that when exposed to *Streptococcus gallolyticus* (*SGG*), a gut pathobiont, the host mucosa promptly adjusts the behavior of specialized goblet cells (GCs) located in the middle of the crypts. A subset of these cells undergoes a transformation, becoming intercrypt goblet cells (icGCs), which do not detach from the surface but instead migrate along intercrypt spaces while secreting mucins. These mucins form a dense layer covering the epithelial cell surface and filling the gaps between mucus plumes secreted from crypt openings, thereby forming a continuous protective mucus layer. Notably, the mucins produced by icGCs exhibit a distinct glycosylation pattern that makes them impermeable to bacterial pathogens. Significantly, a non-piliated *SGG* mutant unable to bind to mucus fail to induce icGCs, allowing its translocation through the mucosa and submucosa. Intriguingly, a closely related mucus-adherent bacterium, *SGM*, which is considered non-pathogenic, also triggers the differentiation of GCs into icGCs. This discovery opens new avenues for treating patients with intestinal diseases characterized by mucus layer deficiencies, such as inflammatory bowel diseases and metabolic disorders. Utilizing mucus-adherent probiotics to induce icGCs represents a promising strategy for reinforcing the mucosal barrier.

**Graphical abstract:** 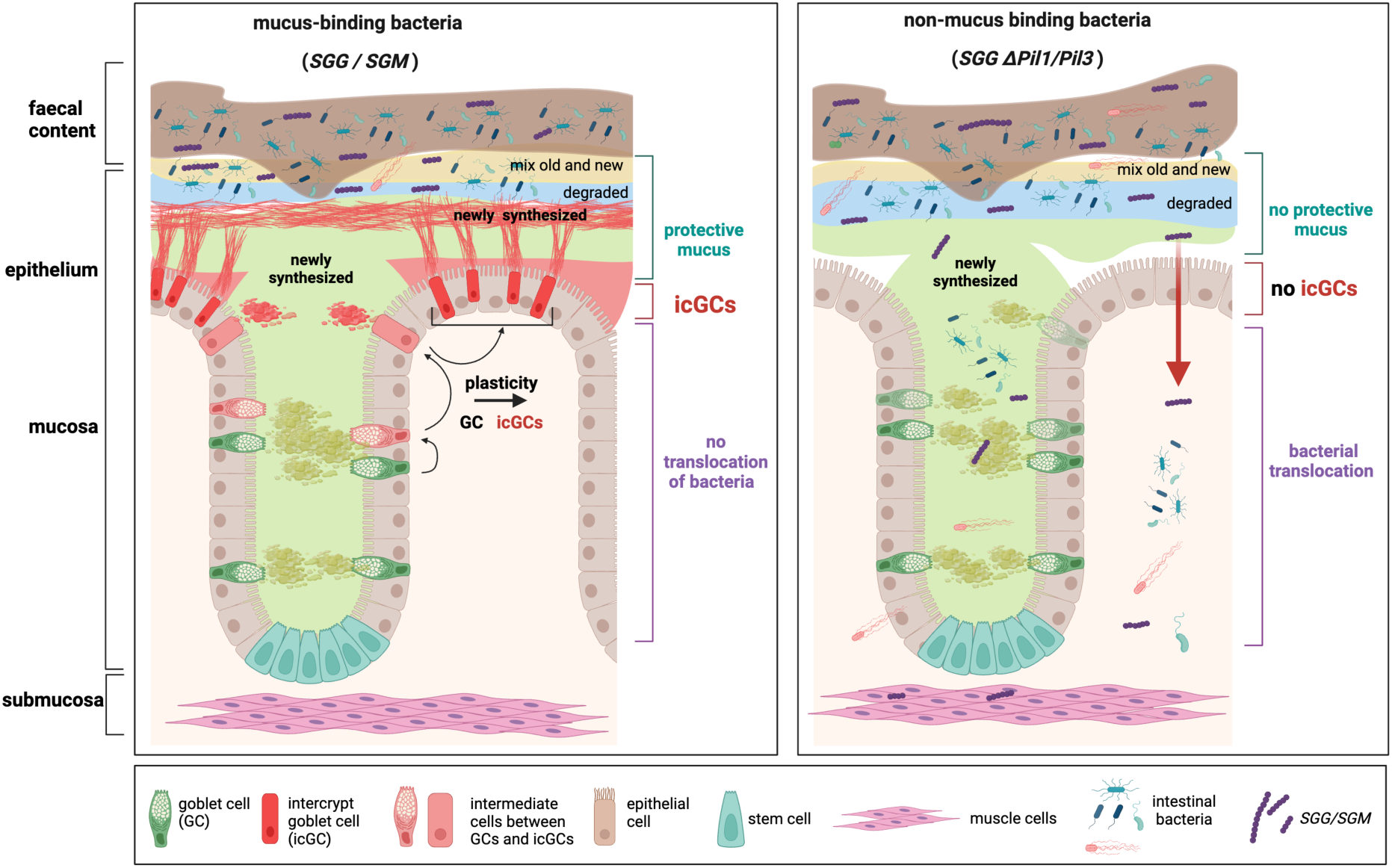

**In brief:** We demonstrate here that, upon oral infection by a gut pathobiont, namely *Streptococcus gallolyticus*, the murine intestinal mucosa displays a novel type of goblet cells recently described as intercrypt goblet cells (icGCs). These icGCs are not shed at the surface of epithelial cells, in contrast to differentiated goblet cells, and produce a continuous protective mucus layer, with a specific pattern of glycosylation rendering it impenetrable to bacteria. No icGCs were induced in response to a non-mucus binding *SGG* mutant, thus allowing bacterial translocation into the mucosa and submucosa, highlighting the essential role played by icGCs in the protective mucus barrier function. Importantly, *SGM*, a commensal mucus-adherent bacterium recognized as safe, is also able to stimulate production of icGCs, opening avenues in the treatment of patients with a “leaky gut”.

**Highlights:**

- Induction of intercrypt goblet cells in murine intestinal tract upon bacterial infection
- icGC potentially arise from differentiated goblet cells through cellular plasticity
- Mucus produced by icGCs is critical for host defense and mucosal barrier function
- Commensal mucus-adherent bacteria also induce icGCs, paving the way for new probiotic treatments for strengthening the mucosal barrier of patients with inflammatory bowel diseases and metabolic disorders

## INTRODUCTION

The human intestine hosts a complex and dynamic microbial ecosystem. In healthy individuals, the gut microbiota maintains a symbiotic relationship with the host, performing crucial functions such as maintaining epithelium homeostasis by stimulating the enteric immune system, digesting food, synthesizing essential nutrients like vitamins, and overall providing a mucosal barrier function^1,2^. This balance is regulated by a constant interaction among three major components of the gut: the microbiota, the mucosal layer, and the epithelial intestinal cells^3,4^. Disruption of this homeostatic balance can lead to various intestinal pathologies including inflammatory bowel diseases, enteric infections, and colorectal cancer^5,6^.

The mucus layer is the first physical barrier separating microorganisms from the underlying epithelium. It is primarily composed of mucins, high molecular weight glycoproteins with numerous O-glycans. In the colon, Muc2 is the main mucin forming a compact inner mucus layer and a detached outer mucus layer that houses the microbiota, including mucus-dwelling bacteria. This outer mucus is loosely attached to the mucosal wall and more firmly bound to feces^7^.

A new model of mucus formation and function in the colon suggests that excreted faeces are covered by two layers of colon mucus: the proximal colon mucus wraps around the faecal mass to seal the microbiota, while the distal mucus produces a second barrier layer around the first^8^. Traditionally, mucins were thought to be secreted by a homogeneous population of goblet cells from the intestinal crypts. However, a recent study has identified several distinct goblet cell populations that form canonical and non-canonical maturation and differentiation trajectories^9^. A specialized surface goblet cell type, the intercrypt goblet cells (icGCs), produces and secretes mucus crucial for forming a protective barrier between bacteria and the epithelium by filling the spaces between mucus plumes secreted by crypt openings. The mechanisms behind the production of icGCs and their potential induction in response to bacterial infections are not yet known. Additionally, while sialylation is known to shape mucus architecture, maintain intestinal host-commensal homeostasis, and inhibit bacterial invasion^10^, little is known about the sialylation of mucus produced by icGCs.

*Streptococcus gallolyticus*, a member of the *Streptococcus bovis*/*S. equinus* complex, includes phenotypically diverse bacteria used in food fermentation, gut commensals and opportunistic pathogens in humans and animals. *S. gallolyticus* is divided into three subspecies, subsp. *gallolyticus* (*SGG*), subsp. *pasteurianus* (*SGP*), and subsp. *macedonicus* (*SGM*). *SGG* is a pathobiont causing septicaemia and endocarditis in elderly and immunocompromised individuals and is often associated with colorectal cancer^11^. *SGM*, the genetically closest *SGG* relative, is considered safe and non-pathogenic and is often used as a control to identify *SGG* virulence traits^12,13^. The first characterized virulence factor in *SGG* was the Pil1 pilus, which mediates binding to collagen and attachment to heart valves, initiating endocarditis^14^. Attachment to colonic mucus and colonization of murine colon depend on another pilus, Pil3^15^. Both Pil1 and Pil3 are heterogeneously expressed in *SGG* through phase variation, a simple regulation mechanism important for avoiding host immune recognition^16^.

To gain deeper insights into how the host mucosa defends against bacterial infection, we investigated the alterations in mucin secretion and glycosylation induced by *SGG* using a murine gut colonization model. We examined whether icGCs could be induced in response to infection and explored the role of sialic acids carried by mucins in host defence against pathogenic bacteria. Our histological observations revealed a massive induction of icGCs upon oral infection with *SGG*. icGC appear to arise from the reprogramming of differentiated goblet cells through cellular plasticity, secreting mucins with specific sialylated O-glycans. Non-pathogenic, mucus-binding *SGM* also stimulated icGCs production, offering a promising approach for strengthening the mucosal barrier in patients with inflammatory bowel diseases and metabolic disorders characterized by increased intestinal permeability.

## RESULTS

### *SGG* induce a significant increase in the number of intercrypt goblet cells *in vivo*

As *SGG* binds to colonic mucins both in vitro and in vivo^15,17^, we hypothesized that *SGG* might alter host mucin production and secretion in mouse model. Antibiotic treated A/J mice were infected orally and sacrificed, and the fixed distal colon were stained by Alcian Blue-Periodic Acid Schiff to detect mucins. Infection with *SGG* WT led to the formation of a high number of intercrypt goblet cells (icGCs). These icGCs were described for the first time in 2021 by Nyström *et al.* as secreting a different mucus as compared to crypt-residing goblet cells (GCs) and constituting an essential component of the intestinal barrier function and protection^9^. icGCs exhibit a typical “rectangular” shape in histological sections after AB-PAS staining rendering them easily distinguishable from “globular” crypt-residing GCs. Compared to the non-infected mice where only a few icGCs were detected (around 35 ± 6 icGCS per mm of epithelial surface, **Fig. 1A**), a significant increase of about 10-fold in the number of icGCs was observed after infection with *SGG* WT with 423 ± 75 icGCs per mm (**Fig. 1B and 1E**). Of note, *SGG* Pil3+, a stronger mucus-binding variant, induced the highest increase in the number of icGCs, with more than 920 ± 89 icGCs per mm (**Fig. 1C and 1E**). Conversely, the non-piliated mutant ΔPil1/Pil3 (deleted for both *pil1* and *pil3* pili) was unable to induce formation of these icGCs (26 ± 6 icGCs per mm; **Fig 1D and 1E**). These results suggest that surface proteinaceous appendages of *SGG*, namely Pil1 and Pil3 pili, previously involved in adhesion to host extracellular proteins, play a key role the induction of icGCs in the murine colon.

**Figure 1.**
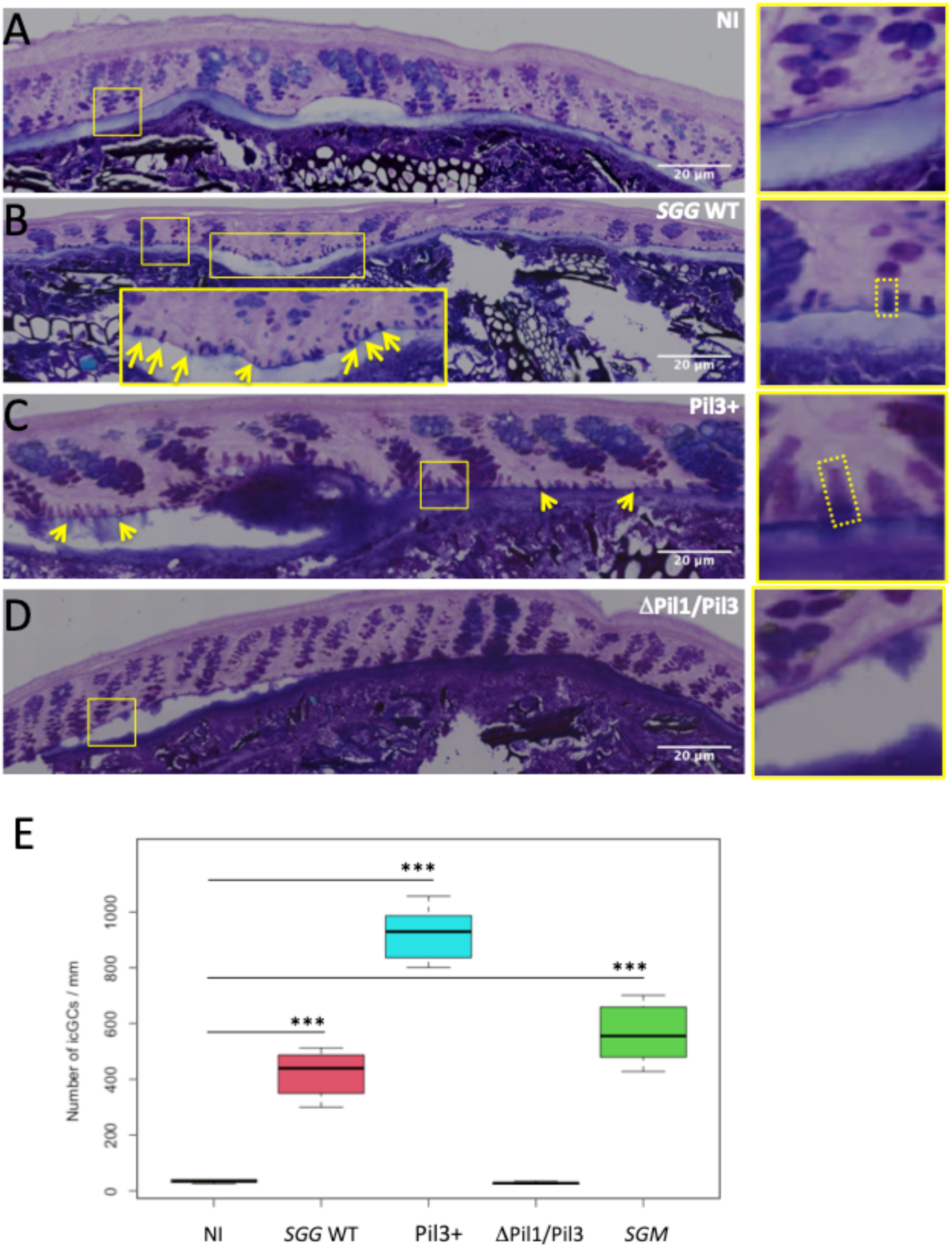
Induction of intercrypt goblet cells (icGCs) after infection of mouse distal colon by *SGG*. Representative distal colonic sections stained with Alcian Blue (AB) and Periodic Acid Schiff (PAS) to visualize acidic and neutral mucins from non-infected A/J mice (A), mice infected orally with *SGG* WT (B), Pil3+ (C) and ΔPil1/Pil3 (D). Only few icGCs are indicated (yellow arrows) but others are easily distinguishable from crypt goblet cells due to their small, typical rectangular shape. Quantification of the number of icGCs per mm, in colon of non-infected mice and mice infected by *SGG* WT, Pil3+ or ΔPil1/Pil3 (E).

### icGCs produce a continuous protective mucus layer at the surface of epithelial cells

Lectins are proteins that recognize and bind specifically to carbohydrate structures within the mucus layer. To investigate icGCs contribution to mucosal layer structure, composition, and function, we used two fluorescently conjugated lectins, UEA1 (recognizing fucose linked in α1,2 to galactose) and SNA (recognizing sialic acids linked in α2,6), in combination with an anti-Muc2 antibody, to stain mucus. In the control non-infected mice, the few icGCs detected were only stained by UEA-1 lectin, as illustrated in the yellow panel of **Fig. 2A**. No staining was observed with SNA and only a weak staining could be observed with the anti Muc2 antibody (**Fig. 2A and 2B**). This agrees with data published by Nylström *et al.* who showed that icGCs were barely detectable using immunostaining of mature Muc2 due to the rapid secretion and reduced storage of mature Muc2 in icGCs compared to crypt-residing GCs. In contrast, mucins from the secreted mucus layer were co-stained with UEA-1, SNA and Muc2.

**Figure 2.**
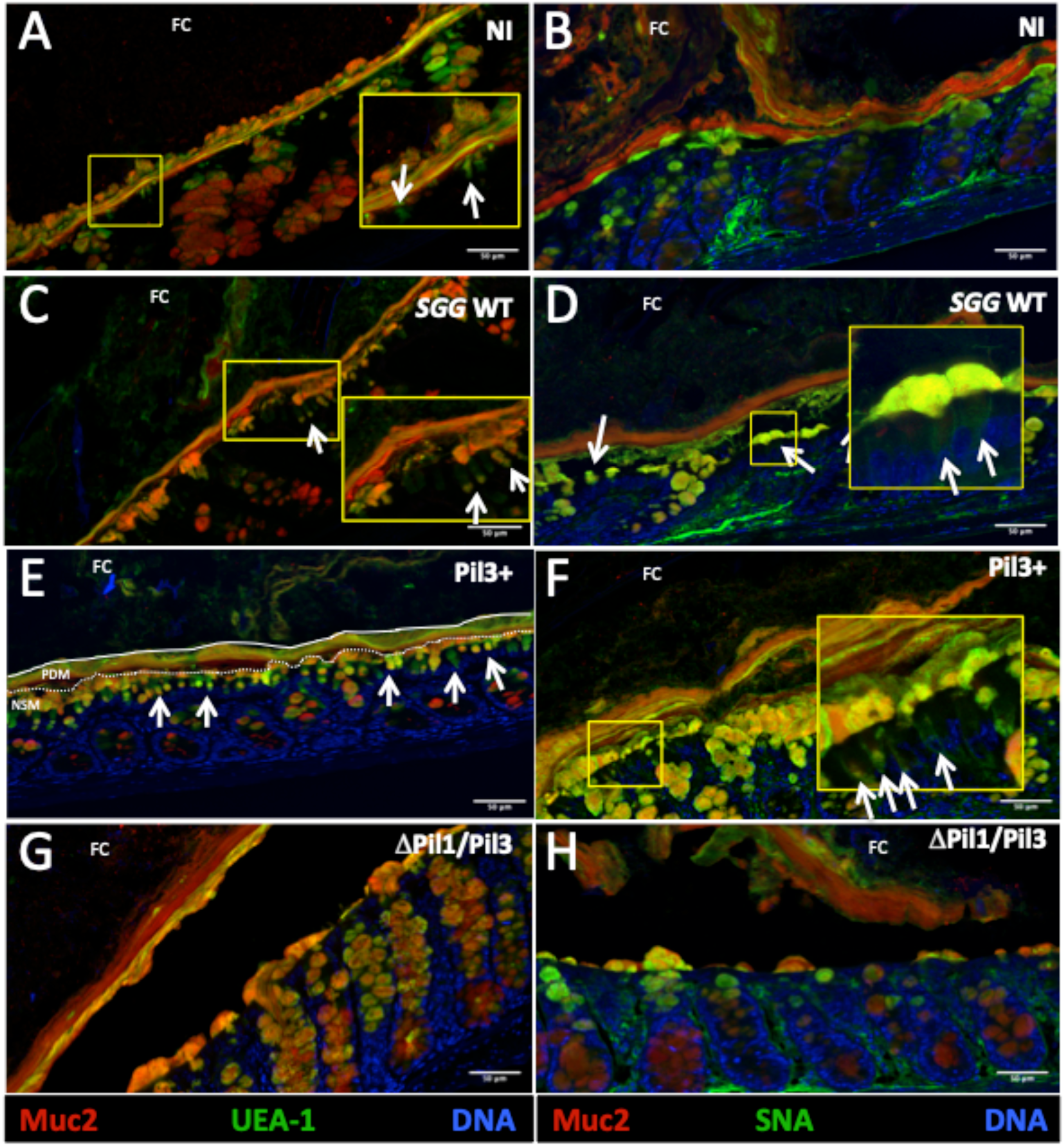
Production of a dense mucus layer by intercrypt goblet cells (icGCs). Representative immunofluorescence staining of mouse distal colonic tissue infected by *SGG*. Colon cross-sections were stained with DAPI to detect DNA (blue), anti Muc2 to visualize colonic mucus (red), and fluorescently conjugated lectins UEA-1 (A, C, E, G) or SNA (B, D, F, H) to visualize fucosylation and sialylation of mucus, respectively (green). Distal colonic sections from non-infected A/J mice (A-B), mice infected orally with *SGG* strains WT (C-D), Pil3+ (E-F) and ΔPil1/Pil3 (G-H). The yellow panels are enlarged views of the boxed regions within the cross-sections. Only few icGCs are indicated (white arrows) but others are easily distinguishable from crypt goblet cells due to their small, typical rectangular shape. FC: faecal content; PDM: partially degraded mucus; NSM: newly synthesized mucus

In mice infected by *SGG* WT, icGCs were barely stained by UEA-1 and SNA (**Fig. 2C and 2D**), and some of them were more intensively co-stained by UEA-1 and Muc2. icGCs secrete filamentous mucins which mix with mucus plumes from crypt-residing GC to form a continuous net-like arrangement at the surface of epithelial cells, thus constituting a first layer of glycosylated mucins bordering a defucosylated and desialylated mucus layer, as evidenced by the thick red layer only stained by Muc2 (**Supp Fig. 1E**). The co-staining of Muc2 with UEA-1 and SNA was confirmed by plotting the intensity profile (**Supp Fig. 1A and 1B**) and intensity profile of SNA and Muc2 (**Supp Fig. 1C and 1D**).

In mice infected by *SGG* Pil3+, the numerous icGCs were co-stained by UEA-1 and Muc2 (**Fig. 2E**). Only a subset of icGCs in the distal colon were also stained by SNA, as shown in **Fig. 2F** and **Supp Fig. 2A and 2B**. These icGCs were found secreting a dense sialylated mucus at the surface of the epithelial cells, filling the space between crypts. Close to the faecal pellet the mucosal layer was costained by UEA-1, SNA and Muc2 and seemed more vaporous (**Fig. 2E and 2F**).

Interestingly, mice infected with the non-piliated ΔPil1/Pil3 mutant, did not induce the formation of icGCs. A thick secreted deglycosylated mucus layer (around 35-40 µm) bordering the faecal pellet (not stained with UEA-1 and SNA) was observed (**Fig. 2G** and **2H**) as seen in the non-infected mice. A second layer, thinner, was composed by a mix of fully and not-fully glycosylated mucins (around 5-10 µm). The mucosal layer secreted at the surface of epithelial cells was mostly produced by crypt GCs and was not covering the entire epithelium. This mucus, sialylated and fucosylated, was significantly thinner (around 6-10 µm), compared to mucus produced in mice infected by *SGG* WT or Pil3+ (around 25-50 µm).

Altogether, these observations suggest that upon bacterial infection, the specific induction of icGCs permits the secretion of a protective layer of newly synthesized mucus with fully glycosylated mucins bordering a degraded mucus layer arising from proximal and middle colon.

### icGCs arise from differentiated goblet cells of the crypts through cellular plasticity

We next questioned the origin of these icGCs. Our histological observations support the idea that icGC originate from crypt GCs. As shown in **Fig.3**, some goblet cells from the middle and the top of the crypts, adopted the “typical rectangular shape of icGC” before migrating to the epithelial surface and occupying intercrypt spaces. Interestingly, whereas goblet cells from the bottom of the crypts were globular, goblet cells from the middle acquired a large rectangular shape and are probably the precursors of icGCs. During their migration towards the apical surface, these cells became thinner probably adopting a shape mimicking those of enterocytes, allowing them to easily integrate the intercrypt spaces (**Fig. 3A and 3B**). Activation of the signaling pathways underlying this cellular plasticity allowing GCs to differentiate into icGCs is specific since non-infected as well as mice infected with a non-piliated ΔPil1/Pil3 mutant were not able to induce this process (**Fig. 3C, 3D and 3E**).

**Figure 3.**
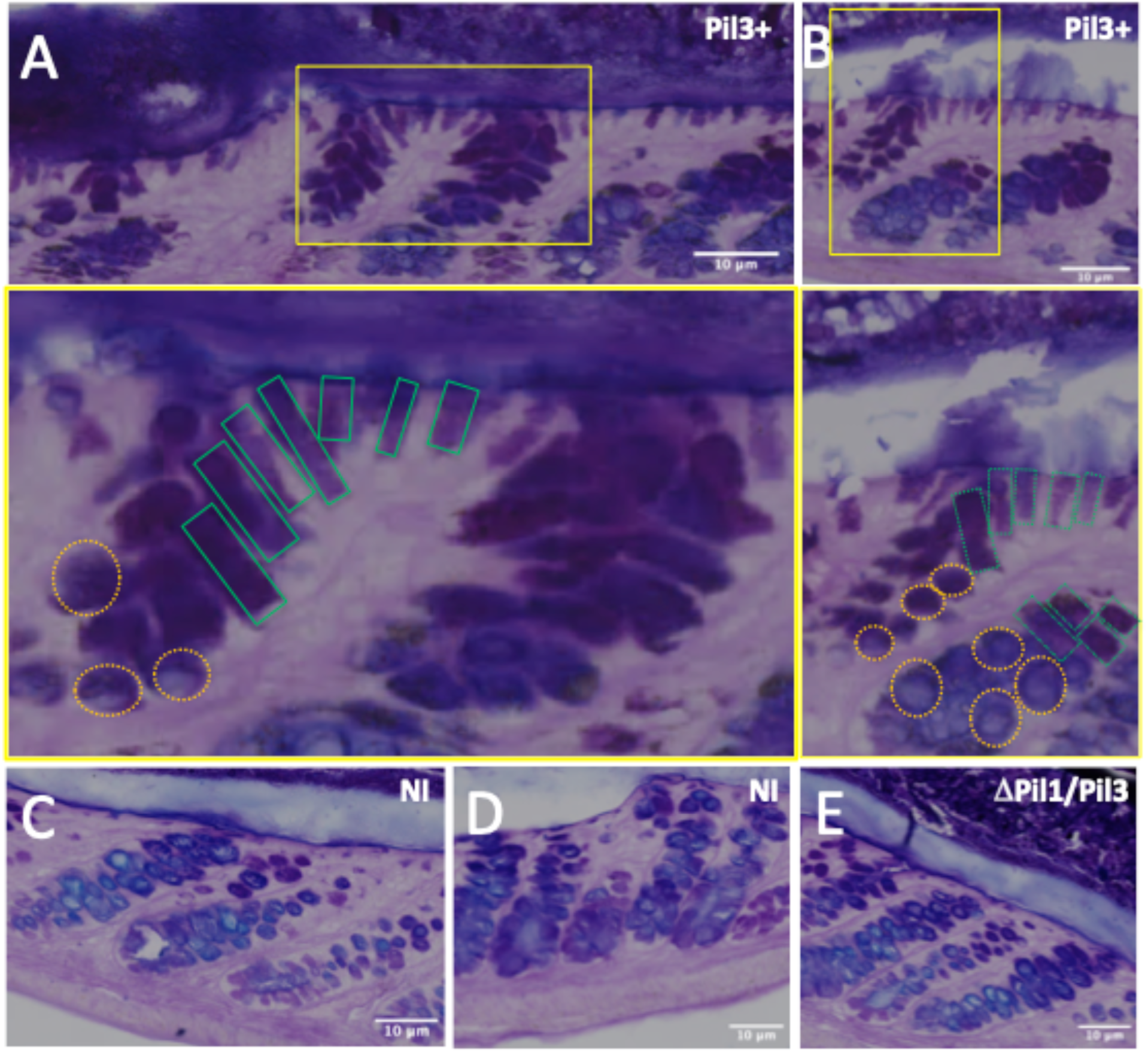
Formation of intercrypt goblet cells (icGCs) from differentiated goblet cells of the crypts. Representative distal colonic sections from mice infected by *SGG* Pil3+ (A, B) or Δpil1/Pil3 (E) and from non-infected A/J mice (C, D) stained with Alcian Blue and Periodic Acid Schiff. The yellow panels are enlarged views of the boxed regions within the cross-sections. The rectangular shape of the icGCs is highlighted by green rectangular boxes whereas the globular shape of GCs is highlighted by orange circles.

### icGCs produce a dense mucus layer impenetrable to bacteria

We next investigated whether the secreted mucus layer could protect against bacterial invasion as a response of the host. Fixed distal colons of mice were stained by SNA lectin and an antibody recognizing *SGG*. In mice infected with *SGG* WT and Pil3+, bacteria were mainly found in the faecal pellets but also in the less dense and less glycosylated mucus layer bordering the faecal material (**Fig. 4A and 4B**). The high density of icGCs produced a continuous and impenetrable layer of mucus keeping bacteria away from the epithelial cells. In *SGG* Pil3+ infected mice, this sialylated mucus was significantly thicker (more than 40 µm) as compared to mice infected with *SGG* WT (around 20-25 µm). None *SGG* could be detected in the mucosa and submucosa. All *SGG* were confined in the faecal pellet, the mucus layer bordering the faeces (composed by a mix a newly synthesized and degraded mucus) as well as the layer of degraded mucus. No *SGG* could be found in the newly synthesized mucus covering the crypt and intercrypt goblet cells (**Fig. 4D**). In sharp contrast, bacterial translocation was clearly seen in mice infected with the non-piliated ΔPil1/Pil3 mutant as attested by the presence of red fluorescence in the mucosa and submucosa (**Fig. 4C and 4D**). These data indicate that the mucus secreted by crypt GC was not able to prevent bacterial translocation in the mucosa and the submucosa. The mucus layer was not continuous at the surface of epithelial cells, leaving many cells unprotected. Even in areas where some icGCs secreted a thin mucus layer, bacterial translocation was noticed, suggesting that this mucus was more porous and penetrable and potentially exhibit a different glycosylation profile as compared to the mucus induced by *SGG* WT and Pil3+.

**Figure 4.**
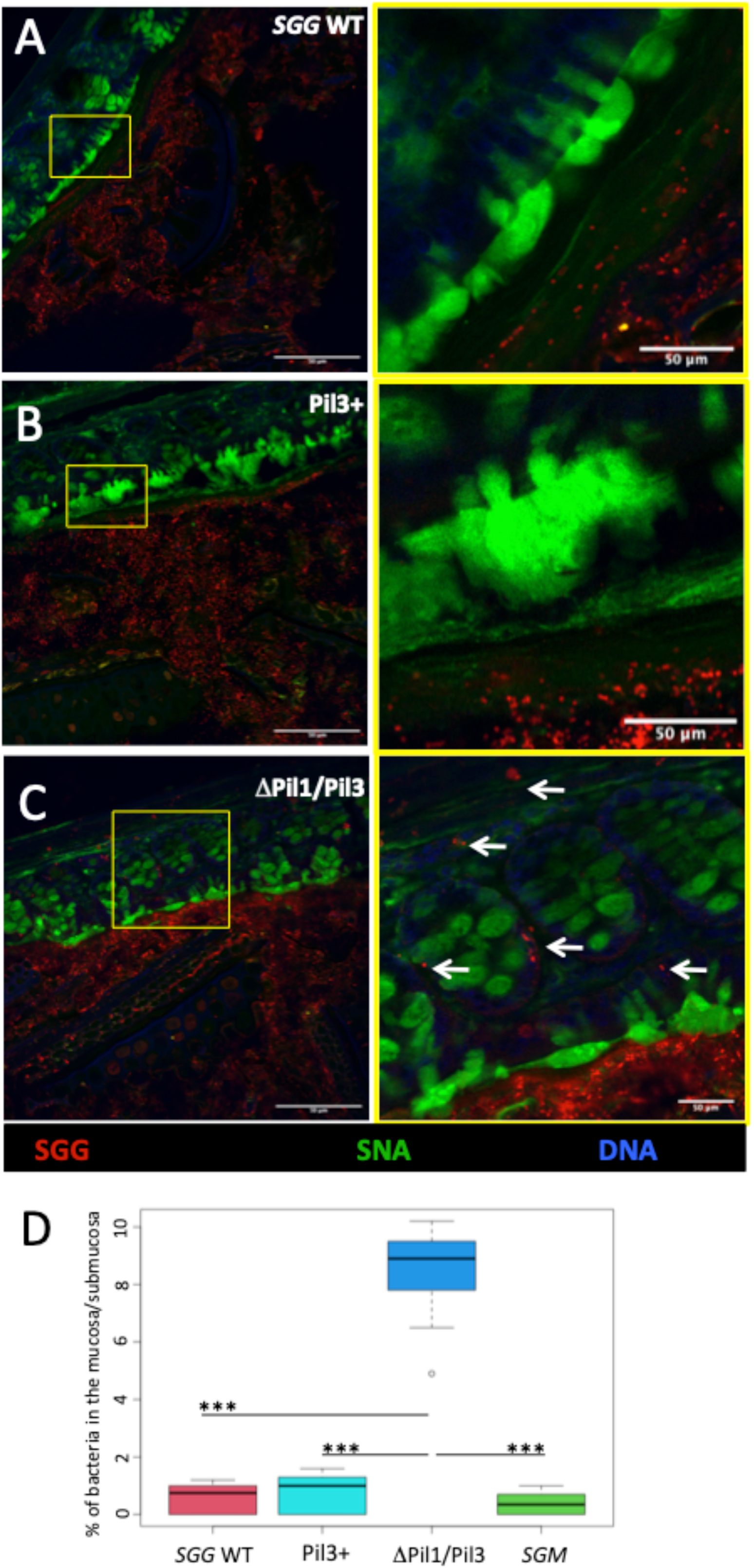
Production of a mucus impenetrable to bacteria by intercrypt goblet cells (icGCs). Representative immunofluorescence staining of mouse distal colonic tissue infected by *SGG*. Colon cross-section was stained with DAPI to detect DNA (blue), anti *SGG* or anti *SGM* (red) to visualize bacteria, and fluorescently FITC labelled SNA lectin to visualize mucus (green). Distal colonic sections from mice infected by *SGG* WT (A), Pil3+ (B) and Δpil1/Pil3 (C) strains of *SGG*. The yellow panels are enlarged views of the boxed regions within the cross-sections. The enlarged view of figure 4C indicates a subpopulation of Δpil1/Pil3 mutant strains that have translocated and are localized in the mucosa and submucosa (white arrows). Quantification of the percentage of bacteria in the mucosa/submucosa compared to the percentage of bacteria in the mucus layer, in the distal colon of mice infected by *SGG* WT, Pil3+ or ΔPil1/Pil3 (E) strains of *SGG* and *SGM*.

### Glycosylation of host colonic mucins is altered upon *SGG* infection

Glycosylation, the process by which sugars are chemically added to proteins, is crucial for the normal function and structure of mucins, which are the main components of mucus. We investigated the glycosylation of mucins in response to *SGG* infection. Altered glycosylation could modify the physical characteristics of mucus, such as viscosity and gel formation, impacting its effectiveness as a barrier against pathogens. To determine whether mucins could be differently glycosylated in response to *SGG* infection, distal colons of mice were collected, mucins were purified and O-glycans were analysed by mass spectrometry. As shown in **Supp Fig. 3A**, a significant increase in the level of mucin sialylation was measured after infection with *SGG* WT and Pil3+ variant up to 41.5% of total oligosaccharides with sialic acid and 27.5% respectively as compared to 13% in control non-infected mice. In contrast, the level of sialylation in mice infected with the ΔPil1/Pil3 mutant was similar to the level in the control group, with less than 20% of O-glycans carrying sialic acids (**Supp Figure 3A**). The structure of sialylated O-glycans were also different between *SGG* WT, Pil3+ groups and NT/ΔPil1/Pil3 mutant groups. Whereas the two main sialylated O-glycans corresponded to ions at m/z 1344 and 1518 for the non-treated mice, the major sialylated O-glycan identified in mucins from *SGG* WT and Pil3+ strains corresponded to an ion at m/z 1589 (**Supp Figure 3B, 3C and 3D**).

### *SGM*, a commensal mucus-adherent bacteria, also induces production of icGCs

To determine if icGCs were only produced after infection by pathogenic bacteria, we next infected mice by *SGM*, a commensal bacterium genetically close to *SGG* but generally recognized as safe^12,13^. *SGM* is a mucus-binding commensal bacterium able to colonize the murine distal colon^17^. Mice infected orally with *SGM* displayed a significant increase in the number of icGCs with 565 ± 104 icGCs per mm compared to the non-treated mice with 35 ± 6 icGCS per mm (**Supp Fig. 4A and Fig. 1E**). Interestingly, *SGM*-induced icGCs were intensively stained by SNA and secreted filamentous mucins (**Supp Fig. 4C**). These icGCs were also stained by UEA-1 but not with mature Muc2 (**Supp Fig. 4B**). As seen previously with *SGG*, icGCS arised from differentiated goblet cells of the crypts through cellular plasticity, as attested by the rectangular shape of goblet cells observed form the middle of the crypts (**Supp. Fig 4D**). The sialylated mucus secreted from icGCs was impenetrable to bacteria (**Fig. 4E**) and was mainly constituted by the O-glycan at m/z 1589, as observed with *SGG* WT and Pil3+ variant (**Supp Fig. 3B, 3C and 3D**).

## DISCUSSION

In this work, we demonstrate for the first time that icGCs are induced in response to bacterial infection, specifically depending on the bacterial ability to bind to the mucus. Specialized intercrypt goblet cells (icGCs) were recently described, with Nystrom *et al.* showing that a deficiency or reduction in icGC numbers leads to an impaired mucosal barrier, as observed in UC patients^9^. While the signaling or environmental cues inducing icGCs formation were not identified, transcriptomic and proteomic profiles of icGCs indicated an increased expression of genes involved in cellular responses to stress and bacteria^9^.

In our literature review, we found a 2013 paper by Gustafsson *et al.* that showed an induction of icGCs in response to infection by *Citrobacter rodentium* (Fig. 7D), compared to non-infected mice (Fig. 7B)^19^. Although icGCs were not described in 2013, the specific rectangular shape of these cells can undoubtedly be attributed to icGCs.

The induction of icGCs appears to correlate with infection by mucus-binding bacteria, as the non-piliated mutant ΔPil1/Pil3 of *SGG* failed to increase icGCs numbers. In contrast, the *SGG* WT and the Pil3+ strains, which overexpress Pil3 (a pilus involved in binding *SGG* to colonic mucus) did induce strongly icGC formation^15^. Similarly, *SGM*, a commensal bacterium that binds to mucus, and *C. rodentium*, a murine pathogen causing colonic crypt hyperplasia, also induced icGCs formation, highlighting the role of bacteria-mucus interaction in this process. The ability of intestinal epithelial cells to revert to a stem cell state is crucial for regenerating the epithelium at sites of mucosal injury, involving the activation of several signaling pathways to promote cellular plasticity and dedifferentiation^20^. In this study, we observed that some differentiated goblet cells from the middle/top of the crypts adopted a rectangular shape, rather than a globular one, migrating to the apical surface to integrate into the intercrypt spaces. This suggests that, in response to injury and mucus-binding bacterial infection, a regenerative response is activated. Instead of dedifferentiating into intestinal stem cells to repopulate crypts after damage, mature goblet cells dedifferentiate to produce icGCs. The emergence of icGCs from differentiated goblet cells indicates remarkable cellular plasticity within the intestinal epithelium, possibly driven by microbial interactions. This adaptive response might be crucial for rapidly reinforcing the mucosal barrier under pathogenic threat. The contrast in icGC induction between strongly mucus-binding and weakly mucus-binding strains (*SGG* WT/Pil3+/*SGM* vs. *SGG* ΔPil1/Pil3) offers intriguing insights into the differential impacts of microbial species on host cellular architecture and immune responses. Our recent global proteomic study in a murine model indicates activation of the integrins pathways in response to *SGG* infection^18^, corroborating the observed cellular plasticity of goblet cells.

This study also highlights the significant impact of *SGG* infection on mucin glycosylation in mice with a strong increase in mucin sialyation by pathogenic *SGG*. In contrast, infection with the non-piliated, non-adherent ΔPil1/Pil3 mutant did not increase sialylation beyond the control level. Additionally, the structure of sialylated O-glycans differed notably between the strains, with the major sialylated O-glycan in *SGG* WT and Pil3+ infections showing a distinct mass/charge ratio compared to non-infected and mutant-infected mice. This indicates that specific bacterial factors are crucial in modifying the glycosylation landscape of host mucins, potentially affecting host-microbial interactions and *SGG* pathogenicity.

These glycosylation changes likely affect the physical properties of mucus, such as viscosity and barrier formation, and biochemical pathways crucial for immune signaling and microbial adhesion. Collectively, these observations highlight sophisticated mucosal defense strategies involving both structural adaptations, like icGC differentiation, and biochemical modifications, such as mucin glycosylation. This dual response fortifies the barrier against immediate bacterial threats and potentially modulates broader immune responses against long-term microbial challenges.

The study’s implications extend to therapeutic strategies, suggesting that manipulating icGC responses or mimicking protective glycosylation patterns could enhance barrier integrity in diseases characterized by mucosal barrier dysfunction, such as inflammatory bowel disease and colorectal cancer. Further research into the signaling pathways and molecular mechanisms driving icGC differentiation and mucin modification in response to microbial stimuli will be crucial for translating these insights into clinical applications.

## SUPPLEMENTAL INFORMATION

## ACKNOWLEDGMENTS

This work was supported by the Institut National contre le Cancer (INCA, Grant PLBIO16-025). The authors are grateful to the Centre National de la Recherche Scientifique (CNRS) and the University of Lille for their recurrent fundings, and the Plateformes Lilloises en Biologie et Santé (PLBS) for help in using fluorescent microscopy.

## AUTHOR CONTRIBUTIONS

Conceptualization, R.L., E.P.K., S.D., and C.R.M.; methodology, R.L., E.P.K., S.D., and C.R.M.; formal analysis, E.P.K., A.M., C.S., S.D., and C.R.M.; investigations, R.L., E.P.K., E.M., B.M., A.M., L.d.M., J.D., C.S., S.D., and C.R.M.; resources, C.S., P.S., S.D., and C.R.M.; writing – original draft, E.P.K., S.D., and C.R.M.; writing – review and editing, R.L., E.P.K., P.S., S.D., and C.R.M.; supervision, S.D., and C.R.M.; project administration, S.D., and C.R.M.; funding acquisition, P.S., S.D., and C.R.M.

## STAR*METHODS

## KEY RESOURCES TABLE

| REAGENT or RESOURCE | SOURCE | IDENTIFIER |
| --- | --- | --- |
| **Antibodies** |  |  |
| Rabbit polyclonal anti-UCN34 | BBPG+ Lab collection | N/A |
| Rabbit polyclonal anti-*SGM* | BBPG+ Lab collection | N/A |
| Mouse monoclonal anti-MUC5AC | Invitrogen | Cat# MA5-12178 |
| Alexa Fluor 546-conjugated donkey anti rabbit IgG | Invitrogen | Cat# A11035 |
| Alexa Fluor 488-conjugated goat anti mouse IgG | Invitrogen | Cat# A11029 |
| Rabbit polyclonal anti-Muc2 (H-300) | SantaCruz | Cat# SC 15334 |
| Sambucus Nigra Lectin (SNA, EBL) Fluorescein (FITC) | Invitrogen | Cat# L32479 |
| UEA-1 Fluorescein (FITC) | Vector Labs | Cat#FL-1061 |
| **Bacterial and virus strains** |  |  |
| *Streptococcus gallolyticus* *SGG* UCN34-WT | Rusniok et al., 2010 | https://pubmed.ncbi.nlm.nih.gov/20139183/ |
| *SGG* UCN34-pil3+ | Martins et al., 2015 | https://pubmed.ncbi.nlm.nih.gov/26014801/ |
| *SGG* UCN34-Δpil1Δpil3 | This study | N/A |
| *Streptococcus gallolyticus* *subsp. Macedonicus* *SGM* | Institut Pasteur Collection | CIP105685T |
| **Chemicals, peptides, and recombinant proteins** |  |  |
| Todd Hewitt Yeast (THY) broth | Millipore | Cat#T1438 |
| DMEM | Dutsher | Cat# L0104-500 |
| fetal calf serum | Biowest | Cat# S1810-500 |
| Phosphate Buffered Saline (PBS) | Euromedex | Cat# ET330 |
| Paraformaldehyde, 4% PBS | Biotum | Cat# 22023 |
| Triton X-100 | Sigma | Cat# X100-100mL |
| Tween-20 | Sigma | Cat# P5827 |
| Prolong Gold Antifade containing DAPI | Thermo | Cat#P36935 |
| vancomycin | Fisher Scientific | Cat# 10379350 |
| neomycin | Sigma | Cat# N1876-25G |
| metronidazole | Sigma | Cat# M3761-5G |
| amphotericin B | Dutsher | Cat# P4030-250mg |
| ampicillin | Sigma | Cat# A9518-5G |
| Periodic acid schiff (PAS) | Alfa Aesar | Cat#J62171 |
| Protein Free blocking Buffer (PFBB) | Thermo Scientific | Cat# 37572 |
| Guanidine chloride | Euromedex | Cat# EU0451 |
| Ethylenediaminetetraacetic acid (EDTA) | Sigma | Cat# E5134 |
| Tween-20 | Sigma | Cat# P5827 |
| Benzamidine | Sigma | Cat# B6506 |
| oN-ethylmaleimide | Sigma | Cat# E3876 |
| Trypsin inhibitor | Sigma | Cat# P9003 |
| Phenylmethanesulfonyl fluoride | Sigma | Cat# P7626 |
| Schiff's reagent | Sigma | Cat# 3952016 |
| Sodium Hydroxide (NaOH) | Sigma | Cat# 6551074 |
| Potassium Borohydride (BH<sub>4</sub>K) | Alfa Aesar | Cat# 32570 |
| Dimethylsulfoxide (DMSO) | Sigma | Cat# 34869 |
| Iodomethane | Sigma | Cat# I8507 |
| Acetic acid | Carlo Erba | Cat# 401421 |
| 2,5-dihydroxybenzoic acid (DHB) |  |  |
| **Critical commercial assays** |  |  |
| Dowex 50 x 2, 200–400 mesh, H<sup>+</sup> form | Carlo Erba | Cat#LL139450B |
| C18 Sep-Pak column (Oasis HLB 6cc- 200mg) | Waters | Cat# WAT106202 |
| **Software and algorithms** |  |  |
| NIS elements AR software |  |  |
| Zen 2.6 software |  |  |
| R (R foundation for Statistical Computing) | https://www.r-project.org | version 4.2.1 |
| Rstudio | http://www.rstudio.com | version 4.2.1 |
| Other |  |  |
| Axioscan Z1 Zeiss slide scanner |  |  |

## RESOURCE AVAILABILITY

### Lead contact

Further information and requests for resources and reagents should be directed to and will be fulfilled by the lead contact, Catherine Robbe Masselot.

### Materials and availability

All unique resources generated in this study may be available from the lead contact under a material transfer agreement.

### Data and code availability

- This paper does not report original code.
- Any additional information required to reanalyze the data reported in this paper is available from the lead contact upon request.

## EXPERIMENTAL MODEL DETAILS

### Murine models and ethics statement

Animals were housed in the Institut Pasteur animal facilities accredited by the French Ministry of Agriculture for performing experiments on live rodents. Work on animals was performed in compliance with French and European regulations on care and protection of laboratory animals (EC Directive 2010/63, French Law 2013-118, February 6th, 2013). All experiments were approved by the Use Ethics Committee of Institut Pasteur #89, registered under the reference dap180064 and were performed in accordance with relevant guidelines and regulations. We declare that these studies are reported in compliance with the ARRIVE guidelines (REF: Kilkenny, C., Browne, W. J., Cuthill, I. C., Emerson, M. & Altman, D. G. Improving bioscience research reporting: The ARRIVE guidelines for reporting animal research. PLoS Biol. 8, e1000412 (2010)). Mice were housed in groups up to 7 animals per cage on poplar chips (SAFE, D0736P00Z) and were fed with irradiated food at 25 kGy (SAFE, #150SP-25). The facility has central air conditioning equipment that maintains a constant temperature of 22 ± 2 °C. Air is renewed at least 20 times per hour in animal rooms. Light is provided with a 14:10-h light: dark cycle (06:30 to 20:30). Mice were kept in polypropylene or polycarbonate cages that comply with European regulations in terms of floor surface per animal. All cages were covered with stainless steel grids and non-woven filter caps.

### Bacterial strains and growth conditions

*Streptococcus gallolyticus subsp. gallolyticus* (or *SGG*) strain UCN34 whose genome is entirely sequenced^21^ was used as reference (*SGG* WT) to isolate the pilus variant *SGG* Pil3+ ^15^ and a mutant deleted for both *pil1* and *pil3* pilus operons (ΔPil1/ Pil3) was constructed as described previously^22^. *Streptococcus gallolyticus subsp. macedonicus* (or *SGM*) strain CIP105683T belongs to the collection of Institut Pasteur and is considered as safe and non-pathogenic^18,23^. It has been sequenced entirely^24^ and its genome is available on NCBI (https://www.ncbi.nlm.nih.gov/bioproject/PRJNA940176).

Both *SGG* and *SGM* were grown at 37°C in Todd Hewitt Yeast (THY) broth in standing filled flasks or on THY agar (Difco Laboratories). Starter cultures before mice oral gavage were prepared by growing strains overnight in 50 ml THY broth. Fresh THY broth was then inoculated with the overnight culture at 1:20 ratio. Exponentially growing bacteria were harvested at 0.5 OD_600_ for the mouse gavage.

## METHOD DETAILS

### Mice colonization experiments

A/J mice were first imported from Jackson Laboratory (USA) and then bred at the Pasteur Institute animal breeding facility in SOPF condition. Eight-week-old female A/J mice were first treated by oral gavage during one week with a broad-spectrum antibiotic mixture including vancomycin (50 µg/g), neomycin (100 µg/g), metronidazole (100 µg/g), amphotericin B (1 µg/g). Additionally, mice were given ampicillin (1g/L) in drinking water for one week and then switched to antibiotic-free water 24 hours prior to bacterial inoculation. Oral gavage of mice was carried out with PBS 1X (NT), *SGM*, *SGG* WT, *SGG*, *SGG* Pil3+, *SGG* ΔPil1/Pil3 using a feeding needle (∼2×10^9^ CFU in 0.2 ml of PBS 1X per mouse) during 3 consecutive days. 24 h post last gavage, all mice were euthanized and colons were removed by cutting from the rectal to the cecal end. For further histological analyses, all colon tissue sections were fixed for 24 h in Carnoy’s solution (60% absolute ethanol, 30% chloroform and 10% glacial acetic acid) for mucus layer protection.

### Immunohistochemistry of mouse tissues and confocal microscopy

Fixed colon fragments were embedded in paraffin and cut in four-µm-thick sections. Sections were deparaffinized in toluene for 5 min, rehydrated with 100%, 95%, and 70% ethanol, followed by distilled H_2_O, and either stained with Periodic Acid Schiff-Alcian Blue (AB-PAS) or antigen retrieved for immunostaining by boiling sections in sodium citrate buffer (pH 6.0) for 20 min. Sections were then incubated in blocking buffer (Protein Free Blocking Buffer, Thermo Scientific) for 2 h. For visualizing *SGG* strains, rabbit polyclonal anti-*SGG* or anti-*SGM* were used and then probed with Alexa-Fluor 546-conjugated donkey anti-rabbit IgG. For detecting mucus, rabbit polyclonal anti-Muc2 (Santa Cruz, H-300) was used and then probed with Alexa-Fluor 546-conjugated donkey anti-rabbit IgG, and FITC conjugated lectins (SNA, recognizing alpha 2,6 linked sialic acids, and UEA-1, recognizing alpha 1,2 linked fucose, Invitrogen) were used to stain glycosylated mucins. Slides were mounted with Prolong Gold Antifade reagent containing DAPI for DNA staining. Slides were then scanned using Axioscan Z1 Zeiss slide scanner (Carl Zeiss Microscopy GmbH) and images were analyzed with the Zen 2.6 software, or viewed on a Nikon A1 R confocal microscope (Nikon Instruments Inc), and images were taken using NIS elements AR software, and further processed using FiJI (Image J)^25^.

### Mucin purification from mice colons

Mice distal colonic tissues were solubilized in an extraction buffer containing 4 M guanidine chloride, 5 mM ethylenediaminetetraacetic acid (EDTA), 10 mM benzamidine, 5 mM oN-ethylmaleimide, 0.1 mg/mL trypsin inhibitor and 1 mM phenylmethanesulfonyl fluoride) for 24h, mucins were purified by density gradient ultracentrifugation at 417,600 g at 15 °C for 72 h (Beckman Coulter LE80 K ultracentrifuge; 70.1 Ti rotor), and identified by Periodic Acid Schiff (PAS) staining after transfer of the different fractions onto a nitrocellulose membrane by slot-blotting.

### O-glycan release, permethylation and MALDI-TOF-MS

The mucin-containing fractions were pooled, dialyzed, lyophilized, and further submitted to β-elimination under reductive conditions (0.1 M NaOH, 1 M BH4Na for 24 h at 45 °C). Borate salts were removed by several co-evaporations with methanol before purification on a cation exchange resin column (Dowex 50 x 2, 200–400 mesh, H^+^ form). Glycans were released as oligosaccharides-alditols, which were then permethylated, in their anhydrous form, in 200 µL dimethylsulfoxide, 300 µL iodomethane and 1 g NaOH, for 2 h, before adding 1 mL acetic acid (5% (v/v)) to stop the reaction. Further purification was carried out on a C18 Sep-Pak column (Oasis HLB, Waters, Milford, MA, USA) in the presence of methanol. Analyses were performed on a MALDI-TOF-TOF geometry mass spectrometer (Analyzer 4800 type, Applied Biosystems/MDS Sciex, Toronto, Canada) equipped with a laser source. Samples were dissolved in a methanol/water solvent (50:50) and coated on a MALDI target in addition with a 2,5-dihydroxybenzoic acid (DHB) matrix at a volume/volume dilution. The relative percent of each oligosaccharide was calculated, based on integration of peaks on MS spectra.

## QUANTIFICATION AND STATISTICAL ANALYSIS

The results are expressed as mean ± standard deviation (SD). Statistical analyses were performed using R (version 4.2.1; R foundation for Statistical Computing) and Rstudio (R: 4.2.1) software. Differences in the level of expression of mucin O-glycosylation were analysed using the Student’s *t*-test. A *P*-value <0.05 was considered statistically significant.

