## Supplemental for "Induction of intercrypt goblet cells upon bacterial infection"

**Supplemental Figure 1. Colonic mucus organization in response to infection by *SGG*.** Representative immunofluorescence staining of mouse distal colonic tissue infected by *SGG* UCN34 WT. Colon cross-sections was stained with DAPI to detect DNA (blue), anti Muc2 to visualize mucus (red), and FITC conjugated lectins UEA-1 (A) or SNA (C) to visualize mucus glycosylation with fucose or sialic acid, respectively (green). Only few icGCs are indicated (white arrows) but others are easily distinguishable from crypt goblet cells due to their small « rectangular » typical shape.

Intensity profile of UEA-1 (B) or SNA (D) and Muc2 in distal colon mucus layer (indicated by a straight yellow line), secreted either by icGCs, crypt goblet cells or mucus bordering fecal content. A defucosylated and desialylated (no staining by UEA-1 and SNA) mucus layer is detected lining the mucus newly secreted by icGCs and crypt openings.

The yellow panel in (E) is enlarged view of the boxed region from (C) and shows the different layers of mucus observed from the epithelium to the faecal content.

**Supplemental Figure 2. Glycosylation pattern of mucus produced by icGCs.** Representative immunofluorescence microscopy of mouse distal colonic tissue infected by *SGG* Pil3+ stained with Muc2 and SNA (A) or with Alcian Blue Periodic Acid Schiff (B). The yellow panels are enlarged views of the boxed regions within the cross-sections. icGCs are indicated by white arrows.

**Supplemental Figure 3. Mucus glycosylation in distal colon is modulated by bacterial infection.** (A) Relative percentage of neutral, sialylated, and sulfated O-glycans carried by mucins purified from distal colon of non-infected A/J mice, mice infected by *SGG* WT, Pil3+, ΔPil1/Pil3, and *SGM.* Data are expressed as mean ± SD; n = 6 per group. Differences in the level of expression of mucin O-glycosylation were analysed using the student's *t*-test. A *P*-value <0.05 was considered statistically significant. The 25$^{th}$, 50$^{th}$, and 75$^{th}$ percentiles and whiskers at 1$^{st}$ quartile -1.5 * IQR (Q1 -1.5 * IQR) and Q3 + 1.5 * IQR based on the level of expression of ions at m/z 1589 (B), 1344 (C), and 1518 (D). Statistically significant associations are marked with an asterisk. IQR, interquartile; Q, quartile. The black panels are proposed structures of O-glycans corresponding to the ions at m/z 1589, 1344 and 1518. Monosaccharide symbols were chosen according to the Consortium for Functional

Glycomics (CFG) nomenclature. Key: fucose (red triangle), GlcNAc (blue square), sialic acid (purple diamond), galactose (yellow circle), and GalNAc (yellow square).

**Supplemental Figure 4. Commensal mucus-adherent bacterium also induces production of icGCs.**

(A) Representative image of distal colonic sections of A/J mice infected with *SGM* type strain stained with Alcian Blue Periodic Acid Schiff to visualize the mucus layer. (B) Representative immunofluorescence staining of mouse distal colonic tissue infected by *SGM*. Colon cross-sections were stained with DAPI to detect DNA (blue), anti Muc2 to visualize mucus (red), and fluorescently conjugated UEA-1 (B) or SNA (C) lectins (green) to visualize mucus fucosylation and sialylation, respectively. Only few icGCs are indicated (white arrows) but others are easily distinguishable from crypt goblet cells due to their small typical rectangular shape. (D) Representative distal colonic section from mice infected by *SGM* stained with AB-PAS showing the differentiation of goblet cells from the middle of the crypts into intercrypt goblet cells. The yellow panels are enlarged view of the boxed regions. The green rectangular boxes reveals formation of icGCs from crypt goblet cells.

(E) Representative immunofluorescence staining of mouse distal colonic tissue infected by *SGM*. Colon cross-section was stained with DAPI to detect DNA (blue), anti *SGM* (red) to visualize bacteria, and fluorescently FITC-labelled UEA-1 or SNA lectin to visualize mucus fucosylation and sialylation (green).

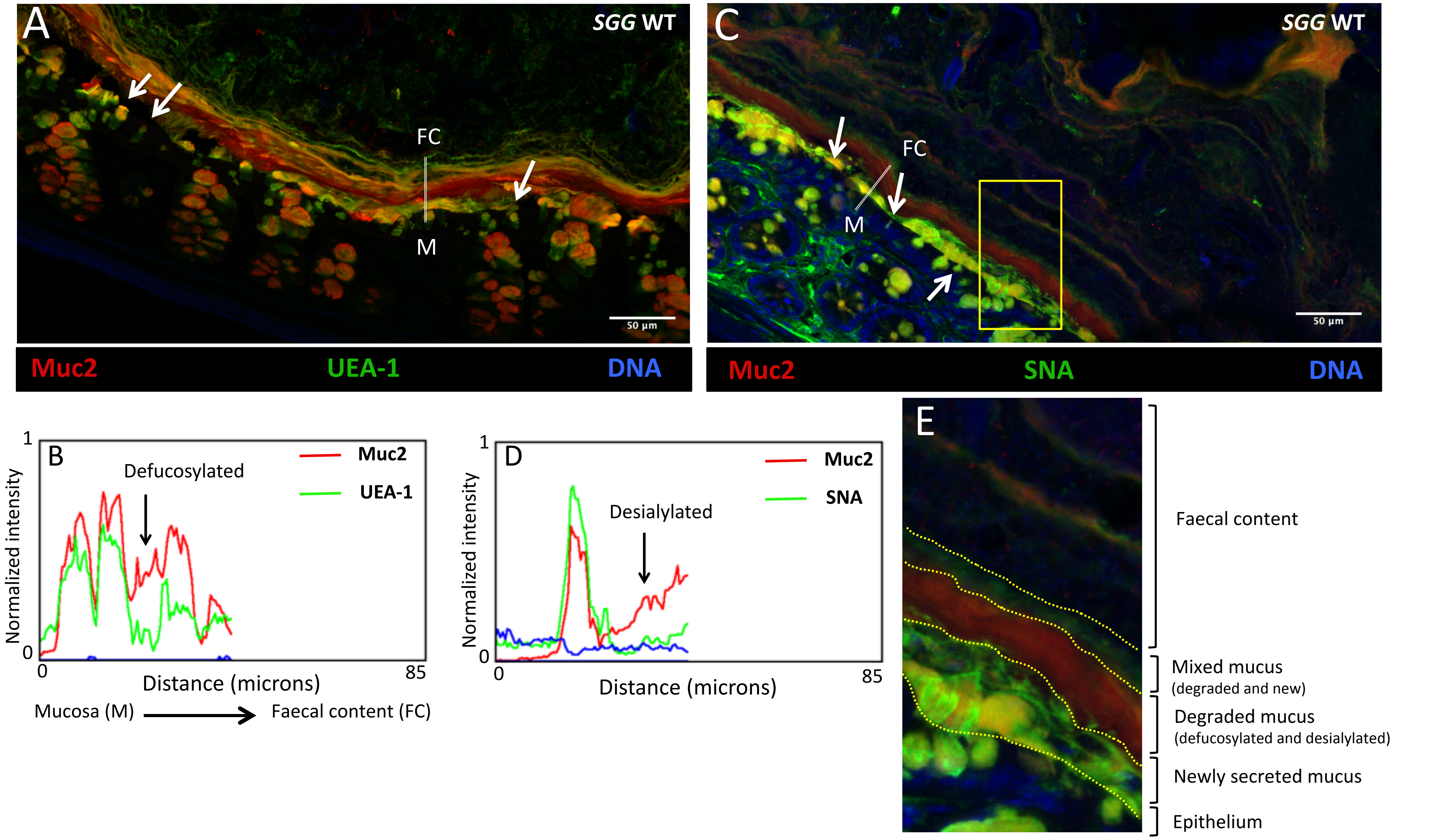
A
SGG WT
FC
M
50 µm
Muc2
UEA-1
DNA
C
SGG WT
FC
M
50 µm
Muc2
SNA
DNA
B
Defucosylated
Muc2
UEA-1
Normalized intensity
1
0
0
85
Distance (microns)
Mucosa (M)
Faecal content (FC)
D
Desialylated
Muc2
SNA
Normalized intensity
1
0
0
85
Distance (microns)
E
Faecal content
Mixed mucus
(degraded and new)
Degraded mucus
(defucosylated and desialylated)
Newly secreted mucus
Epithelium

Fig. S1.

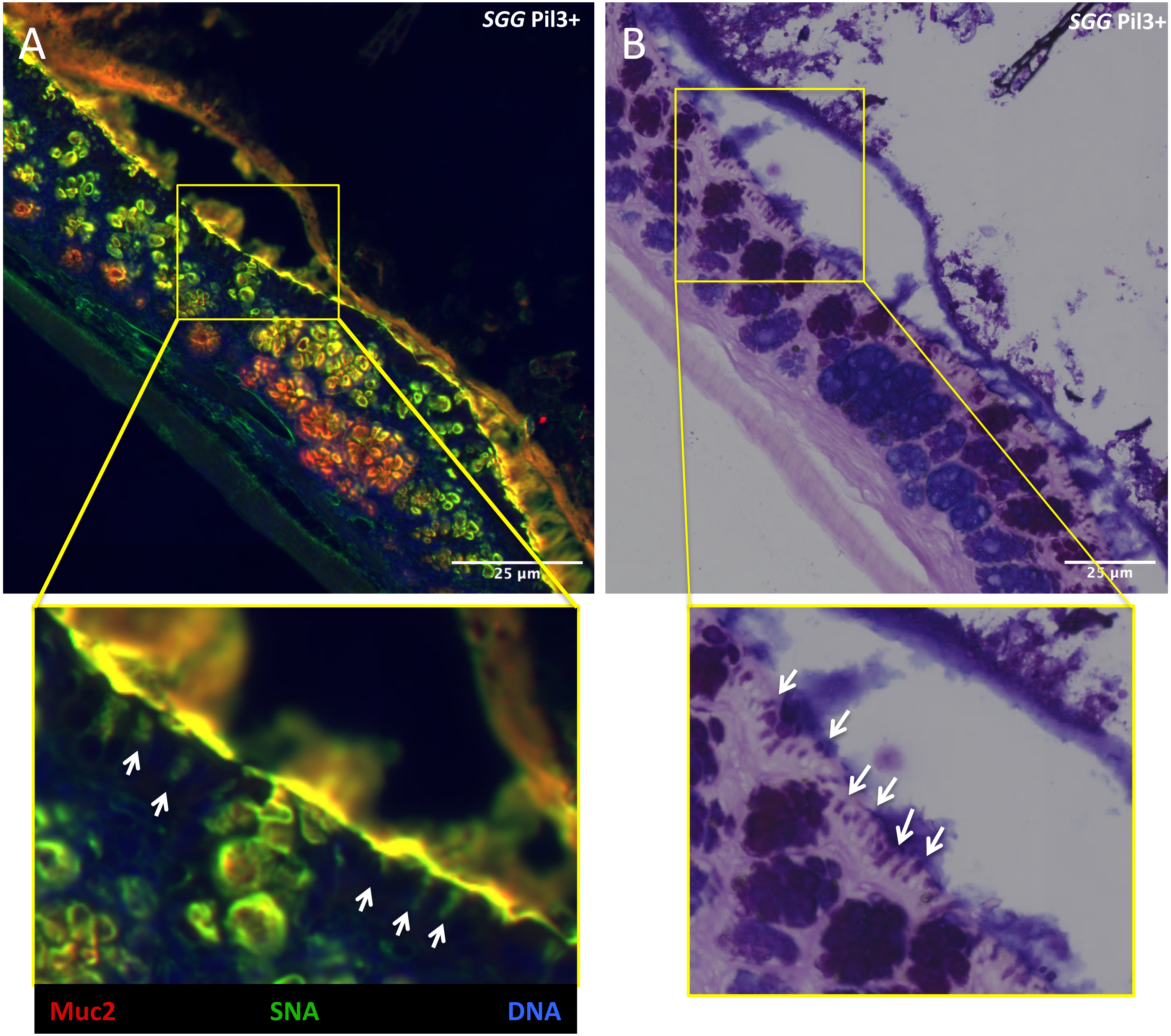


Fig. S2.

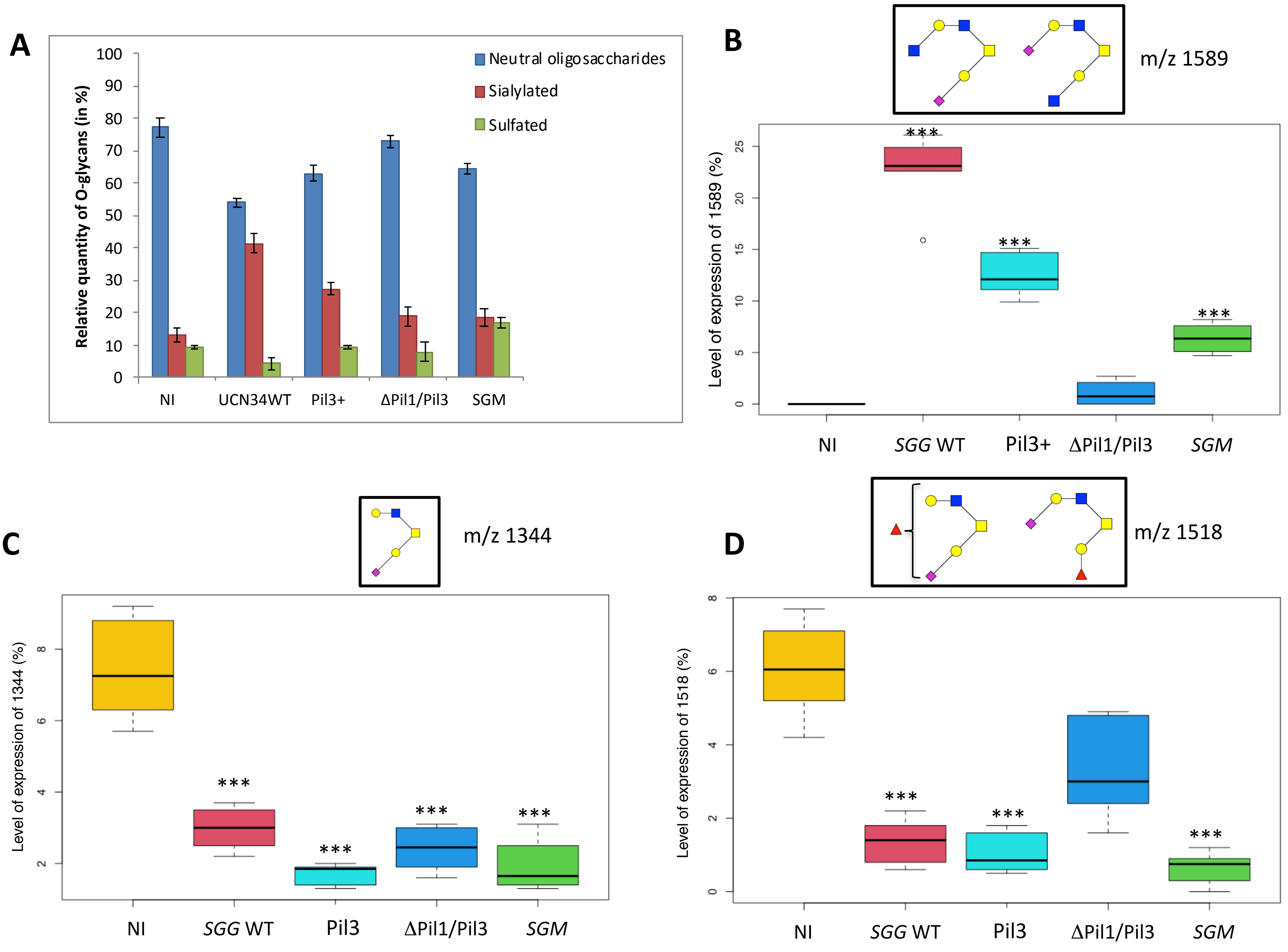

A
Neutral oligosaccharides
Sialylated
Sulfated
Relative quantity of O-glycans (in %)
100
90
80
70
60
50
40
30
20
10
0
NI
UCN34WT
Pil3+
ΔPil1/Pil3
SGM
B
m/z 1589
Level of expression of 1589 (%)
***
25
20
15
10
5
0
NI
SGG WT
Pil3+
ΔPil1/Pil3
SGM
C
m/z 1344
Level of expression of 1344 (%)
8
6
4
2
NI
SGG WT
Pil3
ΔPil1/Pil3
SGM
D
m/z 1518
Level of expression of 1518 (%)
8
6
4
2
0
NI
SGG WT
Pil3
ΔPil1/Pil3
SGM


Fig. S3.

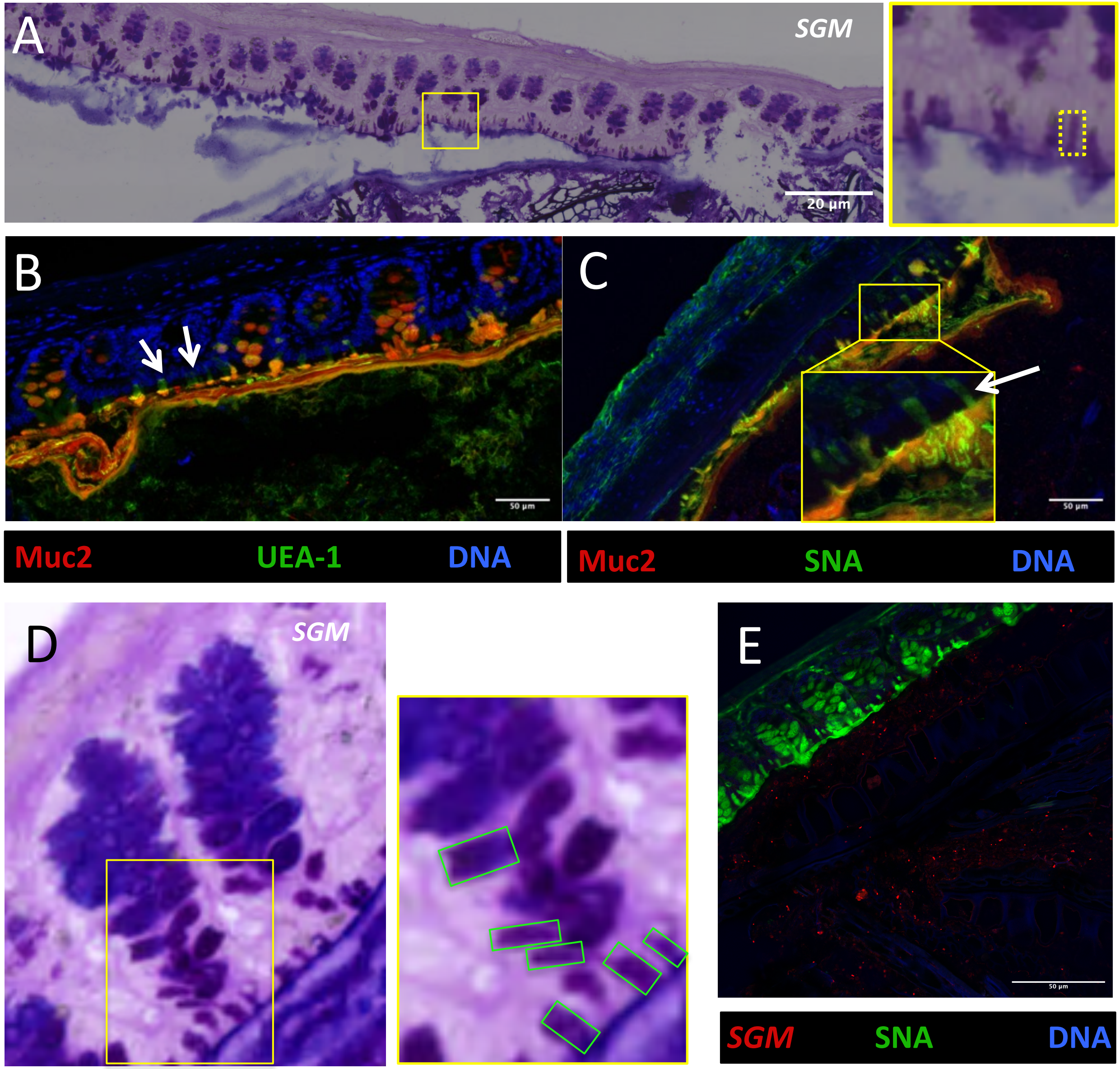
A
SGM
20 µm
B
C
50 µm
50 µm
Muc2
UEA-1
DNA
Muc2
SNA
DNA
D
SGM
E
50 µm
SGM
SNA
DNA

Fig. S4.
